# Osmotically Induced Shape Changes in Membrane Vesicles

**DOI:** 10.64898/2026.04.03.716363

**Authors:** Rajiv G Pereira, Biswaroop Mukherjee, Sanjeev Gautam, Mattiangelo D’Agnese, Subhadip Biswas, Rachel Meeker, Buddhapriya Chakrabarti

**Affiliations:** Department of Physics, Kansas State University, 1228 M. L. K. Jr. Drive, Manhattan, KS, 66506, USA; NewOrbit Space Ltd, 11 S View Park, Marsack St, Reading RG4, UK; Department of Physics and Astronomy, University of Sheffield, Sheffield S3 7RH, UK; Kaplan International College, Palace House, 3 Cathedral Street, London SE1 9DE, UK; Université Paris-Saclay, CNRS, Laboratoire de Physique des Solides, 91405 Orsay, France

## Abstract

We develop a self-consistent free-energy framework in which membrane shape and osmotic pressure are determined simultaneously in a finite reservoir by minimizing bending elasticity and solute entropy. Solute conservation makes osmotic pressure a thermodynamic variable rather than an externally prescribed parameter, producing a nonlinear coupling between membrane mechanics and solvent entropy. This coupling modifies the classical stability condition for spherical vesicles: instability emerges from global free-energy competition rather than the linear Helfrich stability criterion. The resulting critical pressures differ by orders of magnitude from Helfrich predictions and agree with simulations for small and large unilamellar vesicles. The framework is relevant to cellular environments involving biomolecular condensate confinement as well as synthetic vesicles and the development of osmotic-pressure-driven encapsulation platforms.

---

Biological membranes are fundamental structural and functional elements of living systems[1–4], providing compartmentalization[5–7], mechanical integrity[8–10] and regulated exchange with the environment[11–13]. From simple lipid bilayers to complex protein-rich cellular envelopes[14–16], membranes must maintain stability while remaining sufficiently flexible to accommodate growth[17–19], division[1, 20, 21], and environmental perturbations[22–24].

In lower life forms such as bacteria, archaea, and simple eukaryotes, membranes operate under large and fluctuating osmotic gradients that strongly constrain mechanical stability [25, 26]. Osmotic pressure differences across the membrane generate stresses that can dominate shape, tension, and integrity, driving swelling, lysis, and morphological transitions [22, 27–29]. In bacteria, survival under hypo-osmotic shock relies on tension-activated mechanosensitive channels that release osmolytes to prevent rupture [28, 30], while broader osmoregulatory strategies tune turgor through solute accumulation and release [26, 31]. Similar coupling between water activity, membrane mechanics, and volume regulation governs osmoadaptation and morphogenesis in yeast and other eukaryotes, underscoring the need for a quantitative framework linking osmotic forces and membrane elasticity [8, 25, 32]. The Helfrich-Canham free energy [2, 14, 33– 36] has been extensively used to compute the vesicle shapes and shape transformations [37–40]. These models use [37, 41–43] a curvature free energy of the form

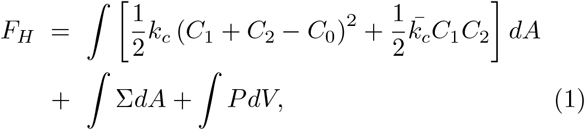

where *C*_1_ and *C*_2_ are the principal curvatures, *C*_0_ is the spontaneous curvature, and *k*_*c*_ and 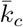 are the bending rigidities associated with mean and Gaussian curvature, respectively. The last two terms impose fixed area and volume constraints via Lagrange multipliers Σ (surface tension) and *P* (pressure difference) [37, 39–41, 43, 44]. For closed vesicles the Gaussian curvature energy integrates to a constant via the Gauss-Bonnet theorem [36, 45] and can be ignored when obtaining the equilibrium shapes via a variational calculation. The problem of determining equilibrium configurations of vesicles has a long history. Deuling et al. [37, 41] categorized axisymmetric vesicle shapes based on the first variation of the Helfrich free energy in Eq. 1. However, the parameterization used, in which the arc length is measured from the north pole *s* = 0 and non-dimensinalized via a scale factor derived from the surface area of a sphere 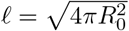 has severe limitations including singularities at the equator *s* = 0.5 and the south pole *s* = 1, rendering integrating the nonlinear shape solutions difficult [46–48]. Instead, a different arclength parameterisation [43, 49–51] removes the singularity at the equator, enabling an accurate determination of shapes. For multivalued membrane configurations other parametrisations [52] have been successfully employed. Irrespective of the model and parametrisation used, the *∫ PdV* term is phenomenologically ascribed to the osmotic pressure acting across the leaflets.

Osmotic pressure has long been recognized as a key control parameter for vesicle morphology within the Helfrich curvature framework, where pressure enters as a Lagrange multiplier enforcing volume constraints [14, 37]. Early theoretical studies showed that increasing pressure destabilizes spherical vesicles and drives shape transitions, including budding and nonaxisymmetric deformations [40, 42]. Subsequent work systematically mapped pressure–curvature phase diagrams for axisymmetric and nonaxisymmetric shapes in spontaneous-curvature and bilayer-couple models [43, 53]. While these approaches successfully classify equilibrium morphologies, osmotic pressure is typically treated phenomenologically rather than derived from solute thermodynamics, motivating recent efforts to couple membrane elasticity directly to osmolyte exchange and osmotic stresses [22, 54–57].

A spherical vesicle minimizes Eq. 1 but becomes unstable beyond a critical pressure 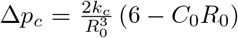, where *R*_0_ is the vesicle radius [37]. Recent experiments on GUVs in hypotonic solutions, however, report critical pressures that exceed this prediction by up to six orders of magnitude [54, 55]. Motivated by this discrepancy, we develop a parameter-free model that couples membrane mechanics to the thermodynamics of solvent exchange. In this framework, osmotic pressure emerges self-consistently from solute concentration differences across the membrane rather than being imposed phenomenologically. We compute equilibrium shapes and phase diagrams, compare them with predictions of purely mechanical bilayer models, and validate our analytical results using coarse-grained molecular dynamics (CGMD) simulations [58, 59] performed via the LAMMPS package [60].

Our starting point is the free energy

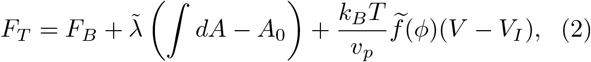

where *F*_*B*_ corresponds to the bending energy term in Eq. 1 (first term), and the second term corresponds to the constant area constraint imposed by the Lagrange multiplier 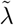 representing the membrane tension. The last term in Eq. 2 describes the solution thermodynamics using a Flory–Huggins free-energy density [61]

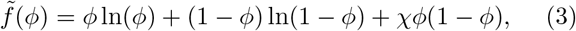

where *v*_*p*_ is the specific volume of the solute and solute volume fraction is

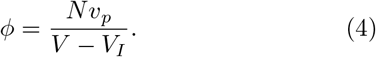

In this expression *V*_*I*_ is the volume of the enclosed vesicle, *N* the number of solute particles, and *V* is the total volume of the reservoir. Since solute particles do not permeate the semipermeable bilayer, we consider an ensemble in which all solute particles remain outside the vesicle. Consequently, Eq. (2) contains no mixing entropy contribution proportional to the vesicle volume. Using a variational formulation, we determine closed axisymmetric vesicle shapes of fixed surface area *A*_0_ that minimize the free energy (2).

Figure 1(c) illustrates the axisymmetric vesicle geometry with symmetry axis *z*. We parameterize the meridional contour by its arc length *s*, measured from the pole *x* = 0 [38, 43], and describe the shape using the radial coordinate *x*(*s*) and the tangent angle *ψ*(*s*) measured relative to the *x* axis. Using arc length parameterization *s*, the relations 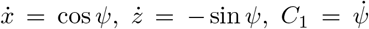, and *C*_2_ = sin *ψ/x* follow, where the derivative is w.r.t. the variable *s*. The enclosed volume of the vesicle is therefore given by *V*_*I*_ = ∫*πx*^2^ sin *ψds*, while *A* = ∫2*πxds* is the surface area. Note that the variables *x, ψ*, and *z* are not all independent and are instead connected via the geometric constraints denoted above. The free energy expression in Eq. 2 thus has an additional term 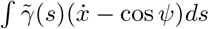 enforcing the geometrical constraint with 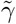 being the Lagrange multiplier. The free energy is thus extremized with respect to *x*(*s*) and *ψ*(*s*) treating them as independent variables. The total free energy is given by 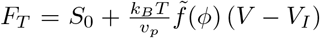, where *S*_0_ is the sum of bending and surface energies expressed in this coordinate system and denoted by,

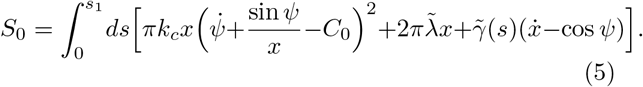

**FIG. 1.**
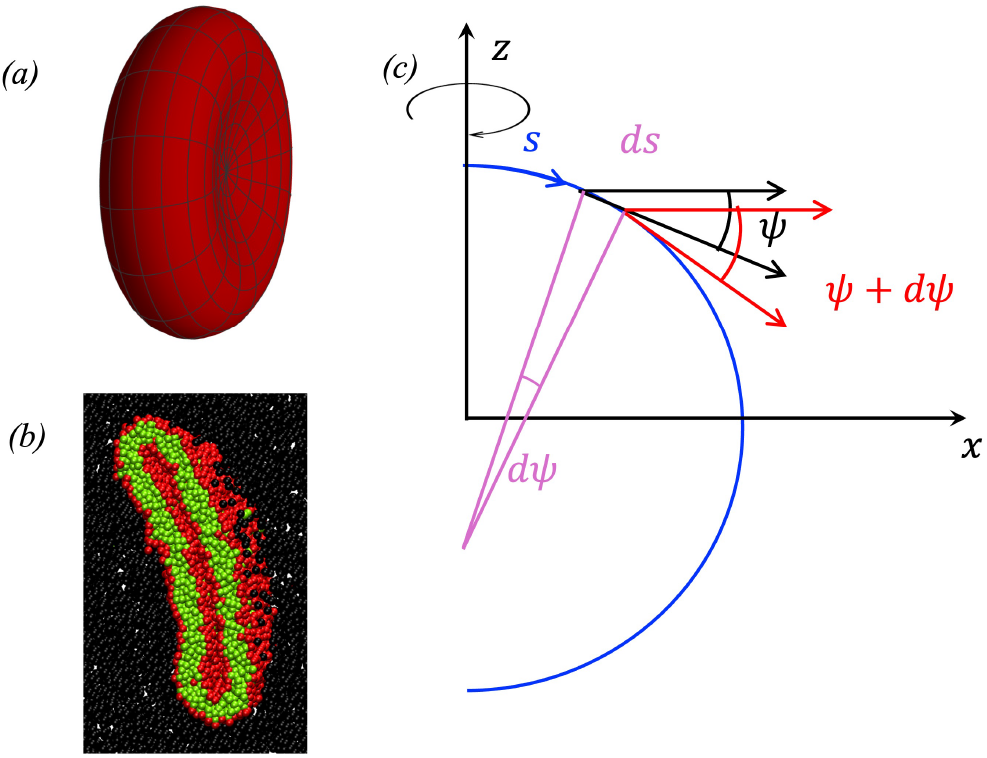
A discotic membrane shape obtained from (a) variational shape calculation, and (b) CGMD simulations of lipid vesicles with hydrophilic head (red) and hydrophobic tail (green) groups in the presence of osmolytes (black). Panel (c) shows an axisymmetric shape in the *z*(*s*), *x*(*s*) plane and the geometric parametrisation used to compute equilibrium vesicle configurations.

Using the expressions for the volume fraction *ϕ* and 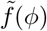 the first variation of the free energy in Eq.(5) is therefore given by *δF*_*T*_ = *δS*_0_ + Π(*ϕ*)*δV*_*I*_, with

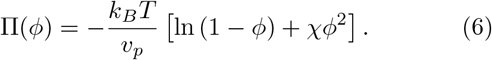

The equations for the contour obtained from the extremization of *F*_*T*_ are

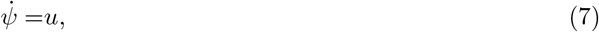

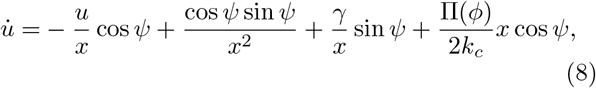

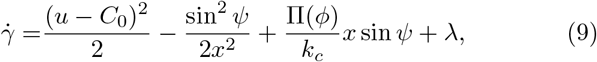

and

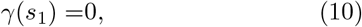

where 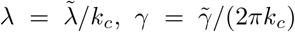, and *s*_1_ is the total arclength. These equations are solved subject to the boundary conditions *x*(0) = 0, *ψ*(0) = 0, *x*(*s*_1_) = 0, and *ψ*(*s*_1_) = *π*, together with the constant area and geometry constraints (see SI). Further, we have *γ*(0) = 0. This is not an independent constraint, and arises from the fact that the variational free energy in Eq.(2) is not an explicit function of the arclength variable *s*, which ensures that the quantity analogous to the Hamiltonian *H* is conserved [38, 51]. Eqs.(7-9) are formally analogous to Seifert’s model [38, 43] with the rescaled pressure term 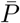 replaced by Π(*ϕ*)*/k*_*c*_ (see Eq. 6). However, this difference is non-trivial and introduces an additional self-consistency condition as Π(*ϕ*), through *ϕ*, implicitly depends on the volume enclosed by the vesicle *V*_*I*_, which is in turn a functional of the shape (*x*(*s*), *ψ*(*s*)).

To obtain self-consistent solutions, we proceed as follows: first, we replace Π(*ϕ*)*/k*_*c*_ with 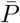 in the shape equations and scale the ensuing equations by a length scale 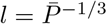 to get the corresponding non-dimensionalised equations. These equations are integrated numerically to obtain the shapes for a given value of *l*^2^*λ* as shown in Fig. 2 that shows five solution branches, namely, sphere, oblate, stomatocyte, and *L*3. Next, we compute the area enclosed by the vesicle in the scaled coordinates (*A*′), and use the constant area constraint, *i*.*e. A* = *l*^2^*A*′ = *A*_0_ to fix the pressure 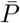, which in turn uniquely fixes the volume of the vesicle 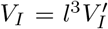, where 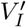 is the volume in the scaled coordinates. We thus have a set 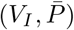 for each solution. These 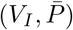 values are plotted together with the function Π(*ϕ*(*V*_*I*_)) for a given value of *N*, with all the other parameters fixed. See inset in Fig. 3 for a plot of Π(*ϕ*) with 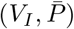 for solutions in the oblate branch. The intersections in this plot represent the solutions that respect the self-consistency condition

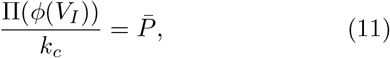

for a given *N*. We compare the total free energy *F*_*T*_ of the self-consistent solutions as a function of the osmolyte number *N*. Figure 3 shows *F*_*T*_ (*N*) for distinct morphologies, revealing successive shape transitions with increasing *N*. A sphere-to-prolate transition occurs at *N* ≃ 1954, followed by a prolate-to-discocyte transition at *N* ≃ 2450. The calculations are performed in a box of volume *V* ≈ 5.1 × 10^4^*σ*^3^, with solute molecular volume *v*_*p*_ ≈ 0.25*σ*^3^ and miscibility parameter *χ* = 2.7 [62]. The bending rigidity is *κ* ≈ 20*k*_*B*_*T*, and the vesicle area is fixed at *A*_0_ ≈ 3300*σ*^2^. The corresponding critical osmotic pressure differences are 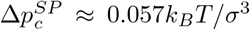 for the sphere–prolate transition and 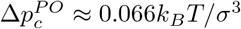 for the prolate–discocyte transition. Further increase in *N* drives discocyte flattening and yields self-intersecting configurations at *N* ≃ 2990, which remain favorable up to *N* ≃ 3161, beyond which the stomatocyte is stable [63].

**FIG. 2.**
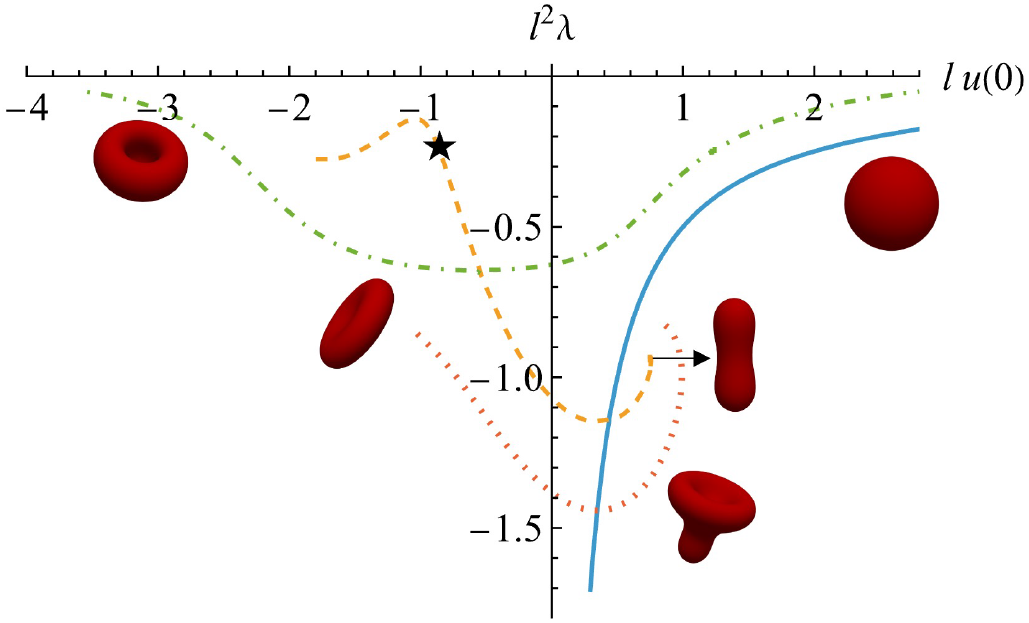
Shape diagram in the *lu*(0) −*l*^2^*λ* plane showing solutions of the scaled shape equations in the absence of osmolytes including (a) spheres (blue solid), (b) stomatocytes (green dash-dotted), (c) prolate and oblate shapes (orange dashed), and (d) *L*3 shapes red dotted. The ⋆ indicates the point on the oblate branch, such that solutions of the shape equations to its left become unphysical, self-intersecting shapes.

**FIG. 3.**
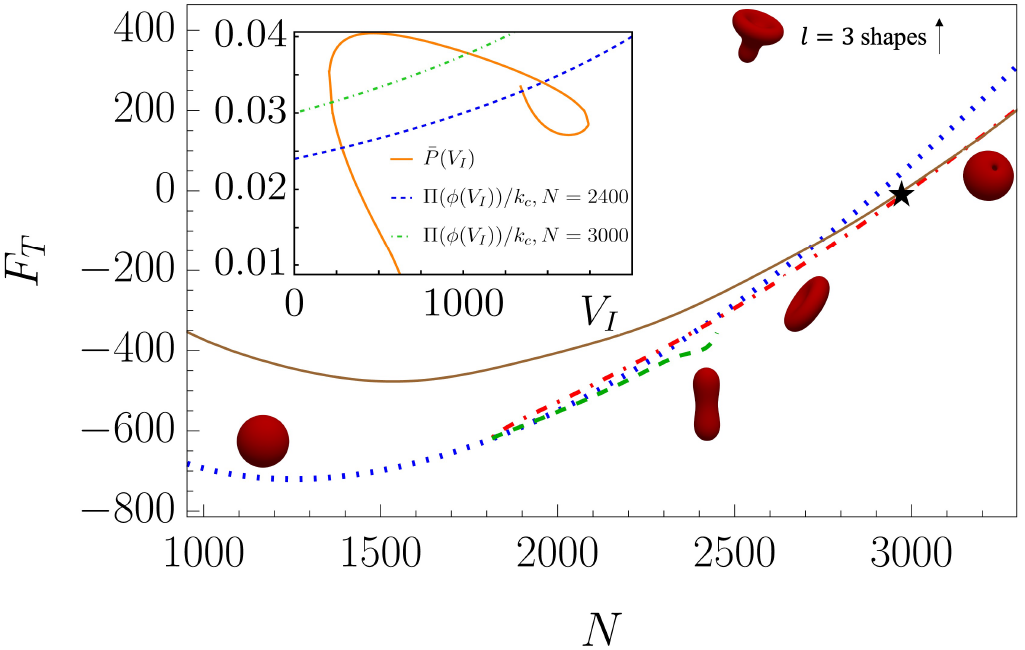
Free energy *F*_*T*_ (in units of *k*_*B*_ *T*) as a function of the osmolyte number *N* for different shapes (a) sphere (blue dots), (b) prolate (green dashed), (c) oblate (red dash dotted), and (d) stomatocyte (brown solid) showing a sequence of shapes that minimize the free energy. The free energy *F*_*T*_ for *L*3 shapes (not shown) are much higher (*>* 1000*k*_*B*_ *T*). The ⋆ indicates values of *N* beyond which self-intersecting shapes appear in the oblate branch. Other self-intersecting solutions of higher energies are omitted. Inset: Shows 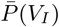 for solutions in the oblate branch (orange solid) and the osmotic pressure Π(*ϕ*(*V*_*I*_))*/k*_*c*_ for *N* = 2400 (blue dashed) and *N* = 3000 (green dash dotted). The intersections correspond to self-consistent solutions of the shape equations.

The self-intersecting solutions are unphysical and reflect the absence of excluded-volume interactions in the Helfrich model; their detailed analysis is beyond the present scope. In contrast, CGMD simulations that incorporate excluded volume yield a flattened discocyte preceding the stomatocyte transition. As shown in Fig. 3 for *N* ≥ 3161, we also observe a transition from a stomatocyte to a double-membrane vesicle. However, complete closure of a stomatocyte into a double-walled vesicle lies outside our framework, as it involves a change in topology that is not captured by the theory. *L*3 shape solutions are observed for *F*_*T*_ ≳ 1000 *k*_*B*_*T* and is not an energy minimum. This high-energy branch compresses the scale and masks the features of interest and is not shown in Fig. 3.

We employ Langevin dynamics simulations to investigate the response of a coarse-grained lipid vesicle subjected to osmotic stress. Membrane vesicles are modeled using the Cooke–Kremer–Deserno model [58, 59] with a three-bead representation of a lipid molecule consisting of one hydrophilic head bead followed by two hydrophobic tail beads performed using the LAMMPS [60] package (see SI). To impose an osmotic imbalance across the membrane, we introduce a third type of coarse-grained particle exterior to the closed vesicle. These particles have diameter *σ* = 1.0 and interact with all membrane beads (both head and tail) exclusively through the repulsive WCA potential, thus ensuring that the osmolytes do not stick or penetrate the membrane, and the concentration differences generate an osmotic pressure acting across the bilayer.

Figure 4(a) shows the phase diagram in the *P*–*V* plane. The pressure is obtained from the virial expression 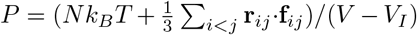, while the volume is computed using a voxel reconstruction approach (see SI). The vesicle volume decreases sharply with increasing osmotic pressure, and representative vesicle shapes are shown in Fig. 4(a). To determine the critical pressure Δ*p*_*c*_ at which the spherical vesicle becomes unstable, we systematically increase the number of osmolytes and compute the relative shape anisotropy (asphericity) parameter (Fig. 4(b)),

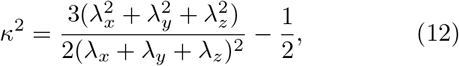

where *λ*_*i*_ are the eigenvalues of the gyration tensor. For the Cooke–Deserno model, we use the parameters *ϵ*_*tt*_ = 1.0 and *ϵ*_*hh*_ = *ϵ*_*ht*_ = 0, with *n*_*ℓ*_ = 8000 lipids forming a vesicle with undeformed outer radius *R*_0_ ≈ 25*σ*. This corresponds to a bending modulus *κ* = 11.97 ± 3.4 *k*_*B*_*T* and a membrane tension Σ = 0.016 ± 0.07 *k*_*B*_*T* calculated from the fluctuation spectrum. The vesicle remains spherical for *N* ≲ 1.5 × 10^4^. As the osmolyte count increases, the vesicle undergoes a sequence of shape transitions: first to a prolate shape for 1.5 × 10^4^ ≲ *N* ≲ 2 × 10^4^, then to a discocyte for 2 × 10^4^ ≲ *N* ≲ 6 × 10^4^, and finally to a stomatocyte for 6 × 10^4^ ≲ *N* ≲ 1 × 10^5^. As the vesicle volume decreases with increasing osmolyte number *N* and osmotic pressure Π(*c*), the asphericity parameter *κ*^2^ increases sharply between *N* = 10^4^ and *N* = 5 × 10^4^. From this transition we estimate a critical osmolyte number *N*_*c*_ ≈ 1.5–1.75 × 10^4^, beyond which the spherical vesicle becomes unstable (see SI). Given a simulation box of size *L* = 100*σ*, the osmotic pressure for *N* ≈ 1.5 × 10^4^ solutes in a volume *V* = 10^6^*σ*^3^ is estimated using van’t Hoff’s law as Π = *k*_*B*_*T* (*N/V*) = 0.015 *k*_*B*_*T/σ*^3^ which shows good agreement with those obtained from our analytical calculations for similar values of the bending modulus. Taking *σ* ≃ 0.8 nm as the lipid radius gives Π ≃ 1.2 × 10^5^ Pa ≈ 1.2 bar. The asphericity parameter shows an interesting feature for higher osmolyte concentrations *i*.*e. N* ≳ 10^5^. Here, the asphericity parameter drops to zero at sufficiently long times. This is due to an instability of the discocyte shape that leads to the formation of double-walled spherical vesicles [64].

**FIG. 4.**
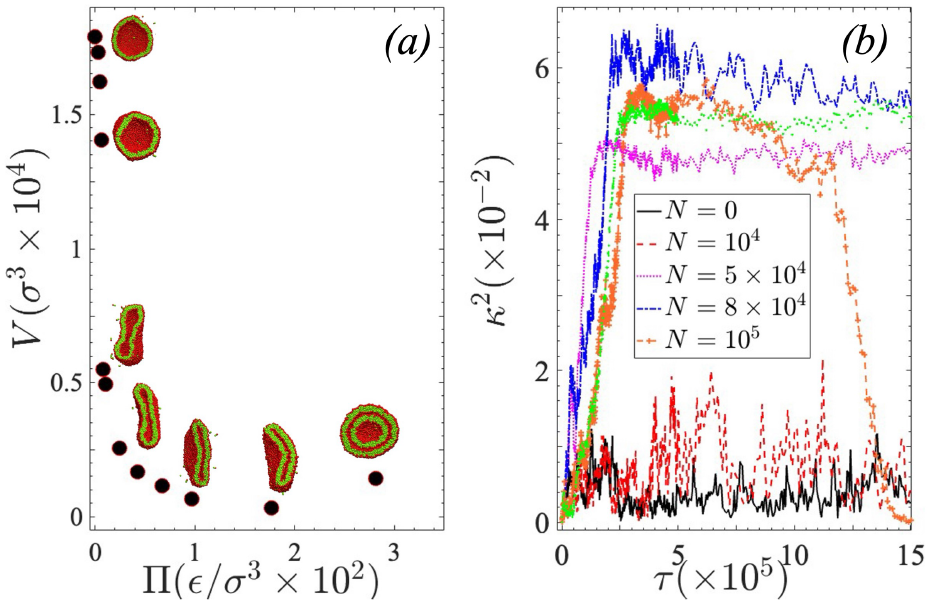
A phase diagram obtained from CGMD simulations in the Π − *V* plane, and the associated equilibrium vesicle shapes (panel (a)). The critical concentration at which a spherical vesicle becomes unstable is identified by calculating the asphericity parameter *κ*^2^ is shown in panel (b), as a function of time *τ* for different osmolyte number *N*. For *N* ≲ 10^4^ *κ*^2^ ≊ 0, and jumps to a finite value once for *N > N*_*c*_ ≈ 1.75 × 10^4^ the mean shape is non-spherical.

Recent experiments on osmotically stressed vesicles [54, 55, 65–67] suggests that the Helfrich–Canham description of membrane elasticity underestimates the critical pressure for vesicle instability by several orders of magnitude, even for large bending rigidities *k*_*c*_ ~ 100*k*_*B*_*T*. There is a fundamental disconnect between the physics operational at different scales, while the elastic theory predicts Δ*p*_*c*_ ~ *k*_*c*_*/R*^3^, for a spherical vesicle of radius *R*, van’t Hoff’s relation Δ*p*_osm_ ~ *k*_*B*_*T* Δ*c* operational at macroscopic scales is independent of the details of the system. It is therefore tempting to compare our results against these experiments. We note that the critical pressure obtained from the Helfrich theory [37, 44] is 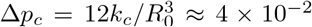 Pa for a GUV of radius *R*_0_ = 5 *µm* assuming *C*_0_ = 0 (*i*.*e*. bilayers), and a bending modulus of *κ* ≈ 97*k*_*B*_*T* [54, 55]. Using van’t Hoff’s relation this yields an equivalent solute number density *N*_*c*_ = Δ*p*_*c*_*/*(*k*_*B*_*T*) ≃ 10^19^ m^−3^ at *T* = 300 K. For CGMD simulations with an osmolyte density *ρ* ≃ 0.015 *σ*^−3^ with *σ* = 0.8 nm, the corresponding number density is *n*_CGMD_ ≃ 3 × 10^25^ m^−3^. We note that extrapolating our analytical and simulation results from SUV/LUV systems to GUVs[54, 55] should be done with caution. The theory presented here connects the shape mechanics with solvent thermodynamics through a nonlinear mechanism. Thus, it is entirely feasible within this theory to have a non-monotonic dependence of the shape transition on the osmolyte concentration, since shape changes are accompanied by volume changes that can be significant in a finite box. This feature is qualitatively different from the classical Helfrich theory where the pressure *P* is an external parameter that increases monotonically, rendering the spherical vesicle unstable beyond a critical threshold.

We present a framework that couples solvent thermodynamics and membrane mechanics to derive a nonlinear instability criterion for spherical vesicles, resolving this discrepancy. Previous work has examined critical osmotic swelling [68, 69], thermodynamic formulations of fluid vesicles [70], and elastic shape phase diagrams in the area– volume ensemble [71]. However, these approaches do not determine osmotic pressure self-consistently from solvent thermodynamics coupled to membrane mechanics, which is the focus of the present work.

Although experiments on osmotically shocked SUV/LUV systems are limited, the magnitude of the effect, governed by the balance between solvent thermodynamics and membrane bending energy, may remain significant in smaller systems near imaging resolution. Such finite-size effects may be particularly relevant when the size of a membrane compartment is only a few times larger than the enclosed object, a situation common in intracellular organelles and condensate–membrane systems. The framework developed here is therefore expected to be relevant to cellular environments, where biomolecular condensates such as the nucleolus or cytoplasmic RNA–protein droplets can mechanically interact with and reshape membrane-bound compartments under confinement and crowding conditions [72–74]. The thermodynamic framework presented here provides a natural starting point for quantitative investigations of such condensate–membrane interactions.

## Author Contributions

Ideation: BC; Conceptualization: BM, RGP, and BC; Analytical Calculations: RGP, BM, MD, RM, and BC; Numerical Simulations: SG, SB, and BC; Drafting: BM, RGP, SG, RM, and BC; Editing: ALL; Funding: BC.

## Acknowledgements

BC thanks the Isaac Newton Institute for Mathematical Sciences, Cambridge, for support and hospitality during the SPL program, where a part of the work on this paper was carried out. This work was supported by EPSRC grant no EP/Z000580/1.

## Notes

### Competing Interest Statement

The authors have declared no competing interest.

