## Supplementary Information for "Osmotically Induced Shape Changes in Membrane Vesicles"

(Dated: April 2, 2026)

In this Supplmenetary Information we outline the derivation of the shape equations of bilayer vesicles deformed by osmotic pressure obtained by the first variation of a modified Helfrich free energy that incorporates solute entropy. We also detail the coarse-grained molecular dynamics simulations following the Cooke-Deserno model of lipid vesicles including the benchmarking protocols used to generate the results reported in the main paper.

### MODEL

The starting point of the outlined method is the Helfrich Hamiltonian[1] that models the phospholipid bilayer bounding the vesicle as a fluid membrane incorporating elastic deformations due to bending.

$$F_H = \int \left[ \frac{1}{2} k_c \left( C_1 + C_2 - C_0 \right)^2 + \frac{1}{2} \bar{k}_c C_1 C_2 \right] dS + \int \Sigma dS + \int P dV. \quad (1)$$

In Eq. 1 $C_1$, and $C_2$ are the two principal curvatures, $k_c$, and $\bar{k}_c$ are the bending moduli, while $\gamma$ is the interfacial tension, and $P = P_{ext} - P_{int}$ is the pressure difference acting across the membrane. The elastic deformation energy contains two terms, one proportional to the mean curvature and the other corresponding to the Gaussian curvature. For closed surfaces, the elastic deformation energy due to the Gaussian curvature integrates to a constant term on account of the Gauss-Bonnet theorem. It is henceforth ignored in the variational calculation of vesicle shapes that are minimizers of the free energy shown in Eq. 1.

A limitation in this framework is that the pressure difference $\Delta p$ is merely ascribed as the osmotic pressure[2] instead of being calculated within the model. This limits the possibility to relate the imposed osmotic pressure to concentration differences acting across the leaflets. To remove this limitation we minimise the total energy of the system composed of the Helfrich free energy in Eq. 1 and a mixing free energy that dictates the partitioning of the solute across the semi-permeable membrane. The total free energy of the system is given by

$$F_T = F_B + F_{mix}, \quad (2)$$

where $F_{mix}$ is the mixing free energy. The mixing free energy is expressed as $F_{mix} = \int \tilde{f}[\phi] dV$, with the free energy density $\tilde{f}[\phi]$ assumed to have the Flory-Huggins form[3]

$$\tilde{f}[\phi] = \frac{F[\phi]}{k_B T} = \phi \ln \phi + (1 - \phi) \ln(1 - \phi) + \chi \phi (1 - \phi). \quad (3)$$

Here, $\phi$ refers to the volume fraction of the solute and $\chi$ the miscibility parameter that dictates the theremodynamic equilibrium between the solute and solvent. The volume fraction is given by

$$\phi = \frac{N v_p}{V - V_I}. \quad (4)$$

where $N$ is the number of solute particles outside the vesicle, $v_p$ the volume occupied by a solute molecule, $V$ the total volume of the system, and $V_I$ the volume enclosed by the vesicle. We assume an initial conformation where the vesicle is impermeable to the solute. The total energy of the vesicle to be minimized is therefore given by

$$F_T = F_B + \frac{k_B T}{v_p} \tilde{f}(\phi)(V - V_I). \quad (5)$$

Our objective is thus to find the closed shape(s) that minimize the free energy (5) for a fixed area $A_0$, restricting ourselves to axisymmetric shapes.

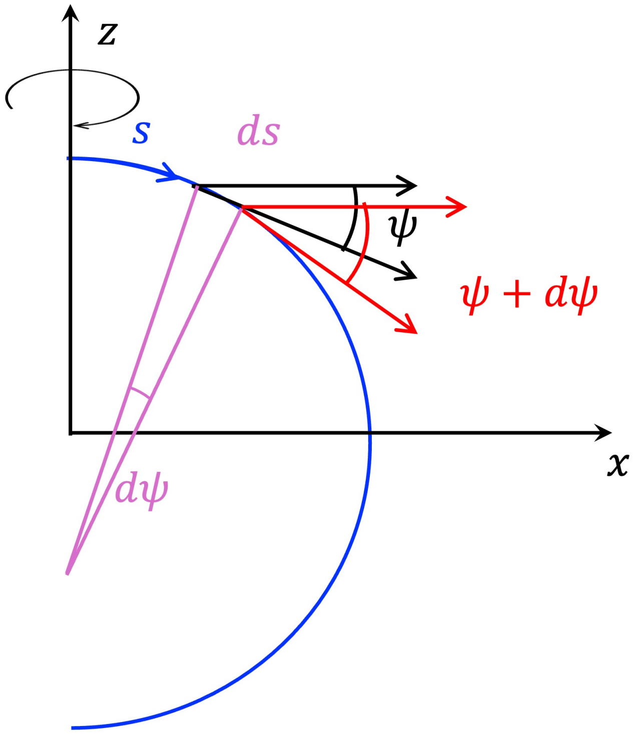


FIG. 1. Schematic of an axisymmetric vesicle shape. The $z$-axis denotes the axis of symmetry, while $s$ is the arclength along the meridional curve $(x(s), z(s))$. The angle $\psi(s)$ between the radial direction and the tangent to the curve varies from 0 to $\pi$ between the two poles.

### GEOMETRIC PARAMETRIZATION

We consider axisymmetric shapes as shown in Fig. 1 with the $z$-axis chosen as the symmetry axis, and $x$ is an axis perpendicular to $z$. We parameterize the contour that defines the edge of the shape in the $x$-$z$ plane using the arclength $s$ along the contour starting from $x = 0$, and use $x(s)$ and the angle $\psi(s)$ substended by the tangent to the contour on the $x$ axis as the coordinates [4]. The following geometric relations can be easily derived.

$$\dot{x} = \cos\psi \tag{6}$$

$$\dot{z} = -\sin\psi \tag{7}$$

$$C_1 = \dot{\psi} \tag{8}$$

$$C_2 = \sin\psi/x, \tag{9}$$

where a dot over the symbols indicate derivative with respect to $s$. The free energy Eq. (5) in these coordinates has the form

$$F_T = \int_0^{s_1} ds\, \pi k_c x \left(\dot{\psi} + \frac{\sin\psi}{x} - C_0\right)^2 + \frac{k_B T}{v_p}\tilde{f}(\phi)\,(V - V_I) \tag{10}$$

with

$$V_I = \int_0^{s_1} ds\, \pi x(s)^2 \sin\psi(s), \tag{11}$$

being the volume of the vesicle. The coordinates $x(s)$ and $\psi(s)$ are not independent but related through Eq. (6). Further, we consider equilibrium shapes having a fixed area $A_0$. Incorporating these constraints into the free energy $F_T$ with the help of Lagrange multipliers $\tilde{\gamma}(s)$ and $\tilde{\lambda}$ yields the quantity

$$F_T = \int_0^{s_1} ds\, \pi k_c x \left(\dot{\psi} + \frac{\sin\psi}{x} - C_0\right)^2 + \frac{k_B T}{v_p}\tilde{f}(\phi)\,(V - V_I) + \int_0^{s_1} ds\, \tilde{\gamma}(s)\,(\dot{x} - \cos\psi) + \tilde{\lambda}\left(2\pi\int_0^{s_1} ds\, x - A_0\right), \tag{12}$$

which we extremize with respect to $x(s)$ and $\psi(s)$ to obtain the equations for the equilibrium shapes. For convenience, we rewrite Eq. (12) as

$$F_T = S_0 + \frac{k_B T}{v_p}\widetilde{f}(\phi)(V - V_I) \tag{13}$$

where

$$S_0 = \int_0^{s_1} ds \left[\pi k_c x \left(\dot{\psi} + \frac{\sin\psi}{x} - C_0\right)^2 + 2\pi\tilde{\lambda} x + \tilde{\gamma}(s)\left(\dot{x} - \cos\psi\right)\right]. \tag{14}$$

Carrying out the first variation we obtain,

$$\delta F_T = \delta S_0 + \frac{k_B T}{v_p}\left[-\widetilde{f}(\phi)\delta V_I + (V - V_I)\delta\widetilde{f}(\phi)\right]. \tag{15}$$

Using the Flory-Huggins free energy Eq. (3) and the expression for $\phi$ Eq.(4) we obtain,

$$\delta\widetilde{f} = \frac{\phi}{V - V_I}\left[\ln\left(\frac{\phi}{1-\phi}\right) + \chi(1 - 2\phi)\right]\delta V_I. \tag{16}$$

Substituting this in Eq. (15) yields

$$\delta F_T = \delta S_0 + \Pi(\phi)\delta V_I \tag{17}$$

where

$$\Pi(\phi) = -\frac{k_B T}{v_p}\left[\ln(1-\phi) + \chi\phi^2\right] \tag{18}$$

is the difference in the osmotic pressures across the membrane. On explicitly carrying out the variation in Eq. (17), subject to the boundary conditions

$$x(0) = 0 \tag{19}$$
$$\psi(0) = 0 \tag{20}$$
$$x(s_1) = 0 \text{ and} \tag{21}$$
$$\psi(s_1) = \pi, \tag{22}$$

we obtain (see Appendix ) the shape equations

$$\dot{\psi} = u \tag{23}$$
$$\dot{u} = -\frac{u}{x}\cos\psi + \frac{\cos\psi\sin\psi}{x^2} + \frac{\gamma}{x}\sin\psi + \frac{\Pi(\phi)}{2k_c}x\cos\psi \tag{24}$$
$$\dot{\gamma} = \frac{(u - C_0)^2}{2} - \frac{\sin^2\psi}{2x^2} + \frac{\Pi(\phi)}{k_c}x\sin\psi + \lambda \tag{25}$$

together with the boundary condition

$$\gamma(s_1) = 0. \tag{26}$$

In Eq. (23) we have used $\lambda \equiv \tilde{\lambda}/k_c$ and $\gamma \equiv \tilde{\gamma}/(2\pi k_c)$. These equations are to be solved with the constraints

$$\dot{x} = \cos\psi \text{ and} \tag{27}$$
$$2\pi\int_0^{s_1} dsxds = A_0. \tag{28}$$

We further have $\gamma(0) = 0$, which is not an additional constraint but automatically satisfied since $F_T$ is not a function of $s$ explicitly (see Appendix ).

If we make the replacement $\Pi(\phi)/k_c \to \bar{P}$, and ignore the dependence on $\phi$, we retrieve the equations in [5], which are

$$\dot{\psi} = u \tag{29}$$

$$\dot{u} = -\frac{u}{x}\cos\psi + \frac{\cos\psi\sin\psi}{x^2} + \frac{\gamma}{x}\sin\psi + \frac{\bar{P}}{2}x\cos\psi \tag{30}$$

$$\dot{\gamma} = \frac{(u-C_0)^2}{2} - \frac{\sin^2\psi}{2x^2} + \bar{P}x\sin\psi + \lambda \tag{31}$$

$$\dot{x} = \cos\psi, \tag{32}$$

with the boundary conditions remaining the same. Note that we can non-dimensionalize these equations by introducing a length scale

$$l = \bar{P}^{-1/3} \tag{33}$$

and defining the dimensionless variables

$$s' = s/l,\ x' = x/l,\ \psi' = \psi,\ u' = lu,\ \gamma' = l\gamma, \tag{34}$$

$$C_0' = lC_0,\ \lambda' = l^2\lambda. \tag{35}$$

In terms of the primed variables Eqs. (29)–(32) reads as

$$\dot{\psi'} = u' \tag{36}$$

$$\dot{u'} = -\frac{u'}{x'}\cos\psi' + \frac{\cos\psi'\sin\psi'}{x'^2} + \frac{\gamma'}{x'}\sin\psi' + \frac{1}{2}x'\cos\psi' \tag{37}$$

$$\dot{\gamma'} = \frac{(u'-C_0')^2}{2} - \frac{\sin^2\psi'}{2x'^2} + x'\sin\psi' + \lambda' \tag{38}$$

$$\dot{x'} = \cos\psi', \tag{39}$$

which is of the same form as Eq. (29)–(32) with $\bar{P} = 1$, thus showing that the shape equations (29)–(32) are scale invariant. On solving these equations for various values of $\lambda'$ we obtain the shape diagram shown in Fig. 2 of main text. The scale invariance of the shape equations is exploited when we fix $l$, and therefore $\bar{P}$, by scaling the solutions meet the area constraint $A = l^2A' = A_0$.

### NOTES ON VARIATION OF $F_T$

Here we demonstrate how the shape equations and the boundary condtions on the geometric factor $\gamma$ are obtained from variation of the total free energy $F_T$. For brevity we introduce the variables $y_1 \equiv x$ and $y_2 \equiv \psi$ and also adopt the convention that the index $i$ is to be summed over whenever repeated within a term. The integrand in Eq. (3) is denoted here by

$$\mathcal{L}(y_i, \dot{y}_i) = \pi k_c y_1\left(\dot{y}_2 + \frac{\sin y_2}{y_1} - C_0\right)^2 + 2\pi\tilde{\lambda}y_1 + \tilde{\gamma}(s)(\dot{y}_1 - \cos y_2). \tag{40}$$

Likewise, the integrand in the expression for volume $V_I$ is denoted by

$$\mathcal{L}'(y_i) = \pi y_1^2 \sin y_2. \tag{41}$$

A smooth axisymmetric shape demands the boundary conditions

$$y_1(0) = 0, \qquad y_2(0) = 0 \tag{42}$$

$$y_1(s_1) = 0, \qquad y_2(s_1) = \pi \tag{43}$$

Suppose the countour $y_i(s)$ extending between $s = 0$ and $s_1$ extremizes $F_T$ over all contours that satisfy the boundary conditions (42) and (43), with the end-point $s_1$ free to vary. Then, any infinitesimal varition about this controur should render $\delta F_T = 0$. Formally, following the standard procedure in variational calculus, we introduce a new contour that is infinitismally close to $y_i(s)$:

$$\hat{y}_i(s) = y_i(s) + \epsilon\eta_i(s), \tag{44}$$

where $\epsilon$ is an infinitesimal parameter. The new contour is assumed to extend from $s = 0$ to

$$\hat{s}_1 = s_1 + \epsilon s_1'. \tag{45}$$

If $\hat{s}_1 > s_1$, then $y_i$ for $s_1 < s < \hat{s}_1$ is defined by Taylor expansion of $y_i$ about $s_1$. Note that the new contour is also forced to satisfy

$$\hat{y}_1(0) = 0, \qquad \hat{y}_2(0) = 0 \tag{46}$$

$$\hat{y}_1(\hat{s}_1) = 0, \qquad \hat{y}_2(\hat{s}_1) = \pi, \tag{47}$$

which ensures that the extremization is done over the subset of curves that respect these boundary conditions. As already stated in the main text, we may break up the variation of the total free energy $F_T$ into two parts as

$$\delta F_T = \delta S_0 + \Pi(\phi)\delta V_I. \tag{48}$$

We first evaluate

$$\delta S_0 \equiv S_0[\hat{y}_i] - S_0[y_i] = \int_0^{\hat{s}_1} ds\, \mathcal{L}(\hat{y}_i, \dot{\hat{y}}_i) - \int_0^{s_1} ds\, \mathcal{L}(y_i, \dot{y}_i). \tag{49}$$

Splitting the first integral in the above immediatly yields

$$\delta S_0 = \int_0^{s_1} ds\, \mathcal{L}(\hat{y}_i, \dot{\hat{y}}_i) + \int_{s_1}^{s_1+\epsilon s_1'} ds\, \mathcal{L}(\hat{y}_i, \dot{\hat{y}}_i) - \int_0^{s_1} ds\, \mathcal{L}(y_i, \dot{y}_i), \tag{50}$$

The first and the last terms are combined to obtain

$$\int_0^{s_1} ds\, \mathcal{L}(\hat{y}_i, \dot{\hat{y}}_i) - \int_0^{s_1} ds\, \mathcal{L}(y_i, \dot{y}_i) = \int_0^{s_1} ds\, \frac{\partial \mathcal{L}}{\partial y_i}\delta y_i + \int_0^{s_1} ds\, \frac{\partial \mathcal{L}}{\partial \dot{y}_i}\delta \dot{y}_i \tag{51}$$

$$= \epsilon \int_0^{s_1} ds\, \frac{\partial \mathcal{L}}{\partial y_i}\eta_i + \epsilon \left.\frac{\partial \mathcal{L}}{\partial \dot{y}_i}\eta_i\right|_{s_1} - \epsilon \int_0^{s_1} ds\, \left(\frac{d}{ds}\frac{\partial \mathcal{L}}{\partial \dot{y}_i}\right)\eta_i, \tag{52}$$

where, in the second line, we have integrated by parts and set $\eta_i(0) = 0$, as we are varying over contours with fixed initial point. Using the above result in Eq. (50) we get

$$\delta S_0 = \epsilon \int_0^{s_1} ds\, \eta_i \left[\frac{\partial \mathcal{L}}{\partial y_i} - \left(\frac{d}{ds}\frac{\partial \mathcal{L}}{\partial \dot{y}_i}\right)\right] + \epsilon \left.\left(\frac{\partial \mathcal{L}}{\partial \dot{y}_i}\eta_i + s_1'\mathcal{L}\right)\right|_{s_1} \tag{53}$$

to $\mathcal{O}(\epsilon)$. From Eq. (44) we obtain

$$\hat{y}_i(\hat{s}_1) - y_i(s_1) = \epsilon\left(\eta_i(s_1) + s_1'\dot{y}_i(s_1)\right) \tag{54}$$

by expanding $y_i$ and $\eta_i$ about $s = s_1$ and retaining terms to $\mathcal{O}(\epsilon)$. Since the left hand side in Eq. (54) evaluates to 0 by virtue of Eqs. (43) and (47), we have $\eta_i(s_1) = -s_1'\dot{y}_i(s_1)$, using which in Eq. (53) immediatly renders it as

$$\delta S_0 = \epsilon \int_0^{s_1} ds\, \eta_i \left[\frac{\partial \mathcal{L}}{\partial y_i} - \left(\frac{d}{ds}\frac{\partial \mathcal{L}}{\partial \dot{y}_i}\right)\right] - s_1'\epsilon \left.\left(\frac{\partial \mathcal{L}}{\partial \dot{y}_i}\dot{y}_i - \mathcal{L}\right)\right|_{s_1} \tag{55}$$

Note that in obtaining this expression for $\delta S_0$ we have not used the explicit form of $\mathcal{L}$ anywhere. Hence, the expression for $\delta V_I$ may be obtained simply by replacing $\mathcal{L}$ in the in Eq. (55) with $\mathcal{L}'$, yielding

$$\delta V_I = \epsilon \int_0^{s_1} ds\, \eta_i \left[\frac{\partial \mathcal{L}'}{\partial y_i} - \left(\frac{d}{ds}\frac{\partial \mathcal{L}'}{\partial \dot{y}_i}\right)\right] - s_1'\epsilon \left.\left(\frac{\partial \mathcal{L}'}{\partial \dot{y}_i}\dot{y}_i - \mathcal{L}'\right)\right|_{s_1}. \tag{56}$$

For generality, we have kept the terms involving the derivative with respect to $\dot{y}_i$ despite $\mathcal{L}'$ not being a function of $\dot{y}_i$. Substituting for $\delta S_0$ and $\delta V_I$ in Eq. (48) using Eqs. (55) and (56), we obtain

$$\begin{aligned}\delta F_T =& \epsilon \int_0^{s_1} ds\, \eta_i \left[ \frac{\partial \mathcal{L}}{\partial y_i} - \left( \frac{d}{ds} \frac{\partial \mathcal{L}}{\partial \dot{y}_i} \right) + \Pi(\phi) \left\{ \frac{\partial \mathcal{L}'}{\partial y_i} - \left( \frac{d}{ds} \frac{\partial \mathcal{L}'}{\partial \dot{y}_i} \right) \right\} \right] \\ & - s_1' \epsilon \left[ \frac{\partial \mathcal{L}}{\partial \dot{y}_i} \dot{y}_i - \mathcal{L} + \Pi(\phi) \left\{ \frac{\partial \mathcal{L}'}{\partial \dot{y}_i} \dot{y}_i - \mathcal{L}' \right\} \right]_{s_1}, \end{aligned} \tag{57}$$

to $\mathcal{O}(\epsilon)$. At the extremal contour, $\delta F_T = 0$ to $\mathcal{O}(\epsilon)$ for any choice of $\eta_i(s)$ and $s_1'$. This implies that the integrand in the first line and the term inside the bracket in the second line evaluated at $s_1$ should both be 0 independently. Further, for the same reason, the integrand in the first line should be 0 for $i = 1$ and 2 independently. Explicitly,

$$\frac{\partial \mathcal{L}}{\partial y_i} - \left( \frac{d}{ds} \frac{\partial \mathcal{L}}{\partial \dot{y}_i} \right) + \Pi(\phi) \left\{ \frac{\partial \mathcal{L}'}{\partial y_i} - \left( \frac{d}{ds} \frac{\partial \mathcal{L}'}{\partial \dot{y}_i} \right) \right\} = 0, \text{ for } i = 1, 2 \tag{58}$$

$$\left[ \frac{\partial \mathcal{L}}{\partial \dot{y}_i} \dot{y}_i - \mathcal{L} + \Pi(\phi) \left\{ \frac{\partial \mathcal{L}'}{\partial \dot{y}_i} \dot{y}_i - \mathcal{L}' \right\} \right]_{s_1} = 0 \tag{59}$$

Equation (58), together with Eqs. (40) and (41), yields the shape equations (9) and (8) in the main text for $i = 1$ and 2, respectively. Likewise, Eq. (59) yields the boundary condition $\gamma(s_1) = 0$.

To capitalize from the fact that neither $\mathcal{L}$ nor $\mathcal{L}'$ is a function of $s$ explicitely, we define the quantity

$$H(s) = \frac{\partial \mathcal{L}}{\partial \dot{y}_i} \dot{y}_i - \mathcal{L} + \Pi(\phi) \left\{ \frac{\partial \mathcal{L}'}{\partial \dot{y}_i} \dot{y}_i - \mathcal{L}' \right\}. \tag{60}$$

which is analogous to the Hamiltonian. It is easily varified that $dH/ds = 0$ along an extremal contour, and $H(s_1) = 0$ by virtue of Eq. (59). It follows that

$$H(0) = \gamma(0) = 0, \tag{61}$$

where the first equality comes from substituting for $\mathcal{L}$ and $\mathcal{L}'$ in Eq. (60) using Eqs. (40) and (41) and then evaluating the ensuing expression at $s = 0$ with the help of the boundary condtions (42).

### CGMD SIMULATIONS

#### CG model of spherical vesicle under osmotic stress

We employ Langevin dynamics simulations to investigate the response of a coarse-grained lipid vesicle subjected to osmotic stress. Membrane vesicles are modeled using a coarse-grained lipid bilayer framework based on the Cooke–Kremer–Deserno model [6, 7]. In this three-bead representation, each lipid molecule consists of one hydrophilic head bead followed by two hydrophobic tail beads.

Non-bonded interactions between coarse-grained beads are modeled using either a purely repulsive Weeks–Chandler–Andersen (WCA) potential or a truncated Lennard–Jones (LJ) potential. The repulsive interaction is given by

$$V_{\mathrm{WCA}}(r) = \begin{cases} 4\epsilon \left[ \left( \frac{\sigma}{r} \right)^{12} - \left( \frac{\sigma}{r} \right)^6 + \frac{1}{4} \right], & r \le r_c, \\ 0, & r > r_c, \end{cases} \tag{62}$$

where $\epsilon$ denotes the interaction strength and the cutoff distance $r_c = 2^{1/6}\sigma$ corresponds to the position of the potential minimum.

The geometric parameters of the lipid beads are chosen as $\sigma_{\mathrm{head,head}} = \sigma_{\mathrm{head,tail}} = 0.95\sigma$ and $\sigma_{\mathrm{tail,tail}} = \sigma$, such that tail beads are slightly larger than head beads. To reproduce the fluid nature of the membrane, additional nonbonded interactions between tail beads are introduced. These effective hydrophobic interactions, which mimic the presence of an implicit solvent, are described by an attractive potential of the form

$$V_{\cos}(r) = \begin{cases} -\epsilon, & r < r_c, \\ -\epsilon \cos^2 \left[ \frac{\pi (r - r_c)}{2 w_c} \right], & r_c \le r \le r_c + w_c, \\ 0, & r > r_c + w_c, \end{cases} \tag{63}$$

where $w_c$ defines the width of the attractive region and the potential smoothly decays to zero over the interval $[r_c, r_c + w_c]$.

To impose an osmotic imbalance across the membrane, additional coarse-grained particles of type 3 are introduced and act as osmolytes. These particles have diameter $\sigma = 1.0$ and interact with all membrane beads (both head and tail) exclusively through the repulsive WCA potential, ensuring excluded-volume interactions without adhesion to or penetration of the membrane. Differences in osmolyte concentration inside and outside the vesicle generate an effective osmotic pressure acting on the membrane.

Bonded interactions between adjacent beads along a lipid molecule are modeled using the finitely extensible non-linear elastic (FENE) potential,

$$V_{\mathrm{FENE}}(r) = -\frac{k r_0^2}{2} \ln\left[1 - \left(\frac{r}{r_0}\right)^2\right], \tag{64}$$

where $k$ is the spring constant and $r_0$ denotes the maximum bond extension.

To enforce a preferred linear conformation of lipid molecules, a bond-angle potential is applied,

$$V_{\mathrm{angle}} = \frac{k_\theta}{2}(\theta - \theta_0)^2, \tag{65}$$

with equilibrium angle $\theta_0 = 180°$ and bending stiffness $k_\theta = 10\epsilon/\sigma^2$.

All molecular dynamics simulations were performed using Large-scale Atomic/Molecular Massively Parallel Simulator (LAMMPS) [8]

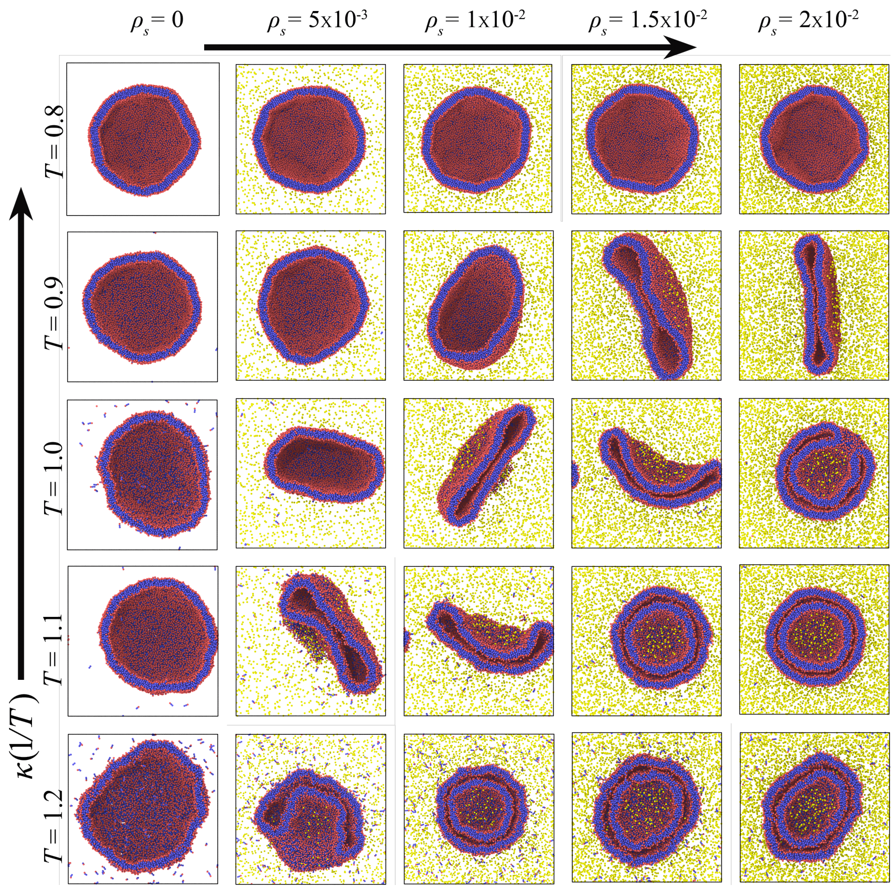


FIG. 2. Temperature-osmotic pressure phase diagram of vesicle deformation in equilibrium. CGMD cross-sectional snapshots of a spherical lipid bilayer vesicle (red being head and blue tail beads) with initial radius $R_0 = 25\sigma$ subjected to changing in external solute (yellow beads) density $\rho_s$ ranging from 0 to $2 \times 10^{-2}\sigma^{-3}$ keeping $w_c = 1.4$ fixed. Temperature is varied from $T = 0.8\varepsilon/k_B$ to $1.2\varepsilon/k_B$, effectively decreasing the bending rigidity. In the absence of $\rho_s$, serve as the reference state, spherical vesicles are mostly undeformed. A similar aspect also emerges at higher membrane bending rigidity or at low temperature. There are deformed and double-vesicle morphologies at higher $T$ and $\rho_s$.

**Osmotic stress - membrane rigidity phase diagram of vesicle deformation:**

To investigate membrane mechanics under osmotic stress, we performed coarse-grained molecular dynamics (CGMD) simulations of lipid bilayer membrane forming an initially spherical vesicle of radius $R_0 = 25\sigma$ with different solute densities $\rho_s$ in Fig. 2. Osmotic pressure in CG simulation is imposed by varying the external solvent particle number density $\rho_s$ from 0 to $2 \times 10^{-2}\sigma^{-3}$, thereby generating a pressure difference across the membrane vesicle interior and exterior. We systematically varied temperatures in the range $T = 0.8\varepsilon/k_B$ to $1.2\varepsilon/k_B$, where increasing temperature effectively reduces the membrane bending modulus $\kappa$. The resulting phase diagram in Fig. 2 reveals that vesicle morphology is governed by the competition between osmotic compression and thermally softened bending rigidity. A similar entropic pressure-driven mechanism of membrane deformation has been observed for a confined polymer under soft membrane confinement [9]. Here, the spherical vesicles largely retain their initial spherical geometry even at finite osmotic stress at low temperature (high bending stiffness $\kappa$). However, as the temperature increases, surface fluctuations reduce $\kappa$ which promotes shape instabilities including prolate, oblate and budded conformations. At sufficiently high osmotic density and high temperature, we observe membrane invagination and the spontaneous formation of double-vesicle (vesicle-within-vesicle) structures. This result demonstrates how thermally tunable bending elasticity controls osmotically driven deformations in lipid bilayer membranes.

**Benchmarked lipid bilayer:**

The coarse-grained lipid membrane model was benchmarked against the Cooke–Deserno reference model [6, 7] to validate its thermodynamic behavior. Systematic variation of the interaction strength $w_c$ and temperature reveals three distinct bilayer phases: gel, fluid, and unstable. The resulting phase diagram is shown in Figure 1.

**Fluctuation Spectrum Analysis**

Vesicle mechanical properties were extracted from production trajectories by computing the spherical-harmonic fluctuation spectrum of the vesicle mid-surface. Head beads (type = 1) were selected and recentered each frame to remove global translation before conversion to spherical coordinates $(r, \theta, \phi)$.

The mid-surface was reconstructed using angular binning on a $N_\theta \times N_\phi$ grid ($24 \times 48$). Within each angular bin, inner and outer leaflet radii were estimated from the radial distribution using percentile statistics (PIN = 15, POUT = 85), and the mid-surface radius was defined as

$$r_{\text{mid}}(\theta, \phi) = \frac{1}{2}\left[r_{\text{in}}(\theta, \phi) + r_{\text{out}}(\theta, \phi)\right]. \tag{66}$$

The reference radius for each frame was taken as

$$R_0 = \langle r_{\text{mid}}(\theta, \phi)\rangle_{\theta,\phi}, \tag{67}$$

and the dimensionless height field was defined as

$$u(\theta, \phi) = \frac{r_{\text{mid}}(\theta, \phi) - R_0}{R_0}. \tag{68}$$

The fluctuation field was expanded in spherical harmonics $Y_{lm}$, and the $m$-averaged spectrum was computed as

$$\langle |u_l|^2\rangle = \frac{1}{2l+1}\sum_{m=-l}^{l} |a_{lm}|^2, \tag{69}$$

with averaging performed over all analyzed frames.

For $l \geq 2$, the spectrum was fit to the continuum quasi-spherical membrane theory including bending and tension contributions, with an additive noise floor $C$:

$$\langle |u_l|^2\rangle = \frac{k_B T}{\kappa(l-1)l(l+1)(l+2) + (\Sigma R_0^2)\, l(l+1)} + C. \tag{70}$$

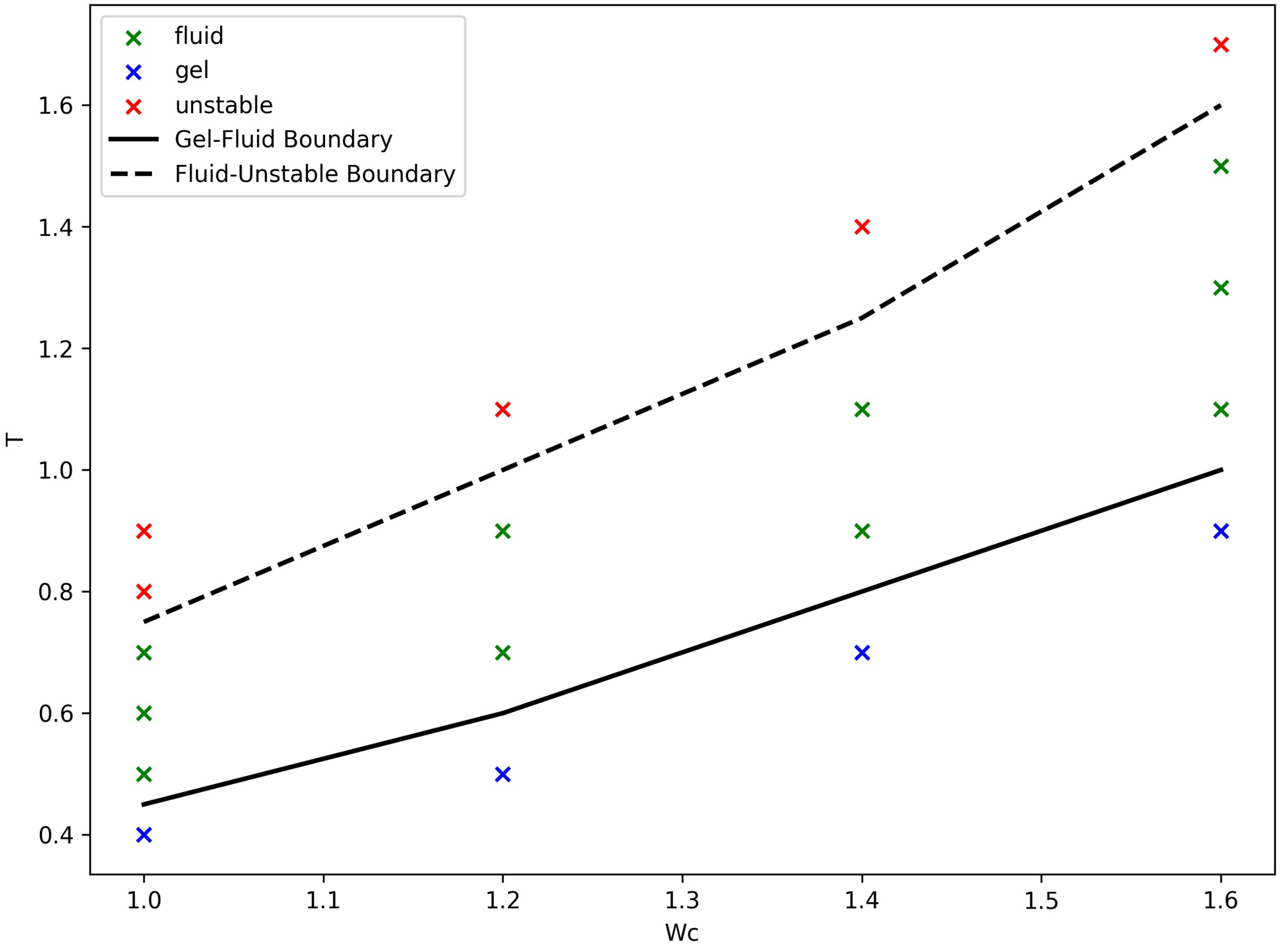


FIG. 3. Phase diagram of the coarse-grained membrane model benchmarking the LAMMPS simulations against the Cooke-Deserno model [6, 7].

To improve parameter robustness, a two-step fitting protocol was employed. First, $\kappa$ and $C$ were determined from a bending-dominated high-$l$ window ($l = 6$–$20$) with $\Sigma R_0^2$ fixed to zero. Second, $\Sigma R_0^2$ was obtained from low-$l$ modes ($l = 2$–$6$) while holding $\kappa$ and $C$ fixed. The surface tension was then calculated as

$$\Sigma = \frac{\Sigma R_0^2}{R_0^2}, \tag{71}$$

with $R_0$ averaged over frames. Parameter uncertainties were obtained from the covariance matrices of nonlinear least-squares fits. The resulting fluctuation spectrum and two-step fit are shown in Fig. 4.

**Asphericity Analysis**

Vesicle shape anisotropy was quantified using the relative shape anisotropy $\kappa^2$ computed from equilibration and production trajectories, which were combined into a single continuous time series. Tail beads were used to construct the vesicle surface. If the production run restarted its timestep counter, timesteps were shifted to ensure continuity.

For each frame, a mid-surface point cloud was constructed using angular binning on a $N_\theta \times N_\phi$ grid ($30 \times 60$). Particle positions were recentered, converted to spherical coordinates, and classified into inner and outer leaflets based on the median radial distance. Within each angular bin, leaflet-resolved radii and directions were averaged and

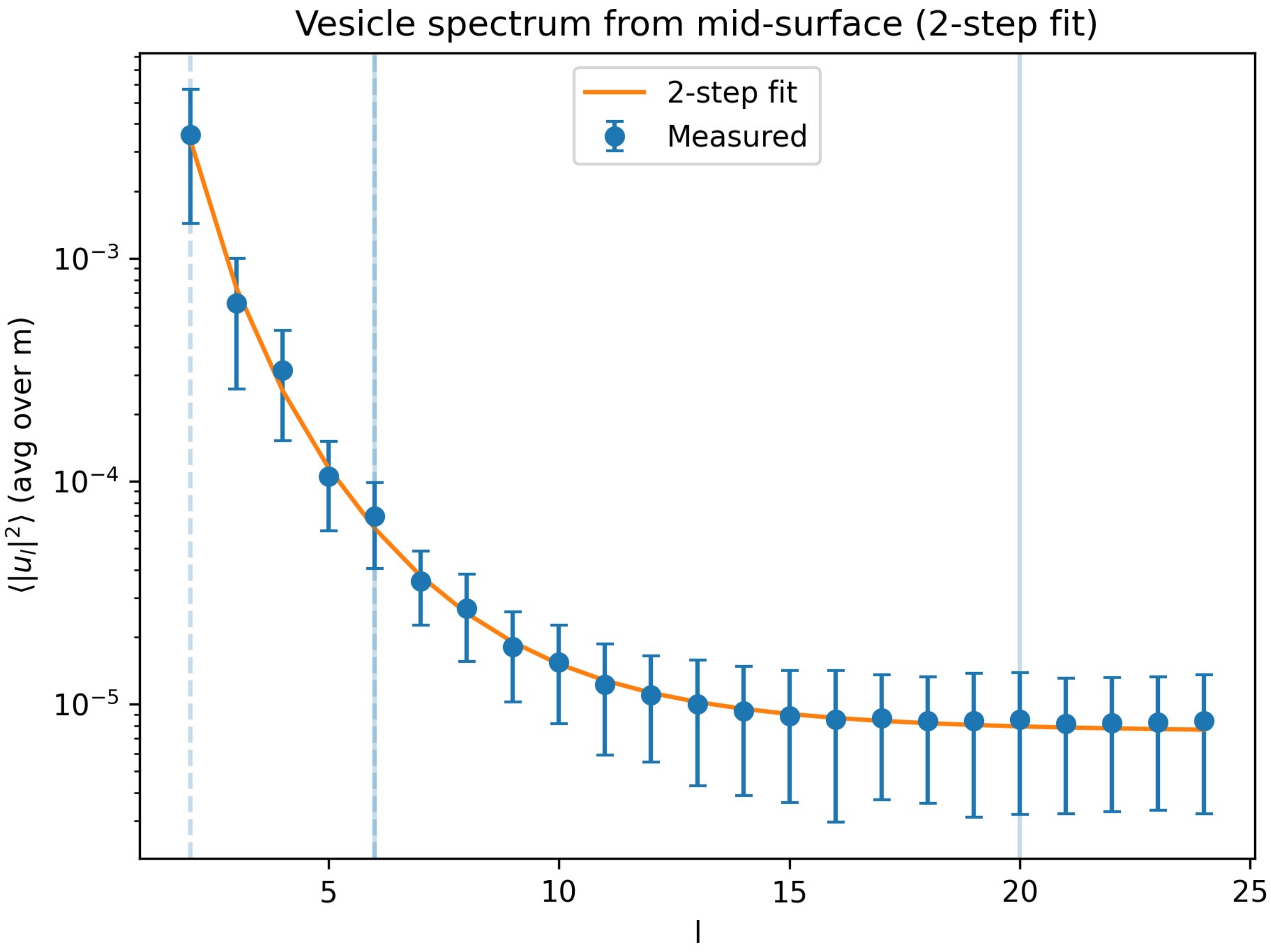


FIG. 4. Spherical-harmonic fluctuation spectrum of the vesicle mid-surface. Points represent the measured mode amplitudes $\langle |u_l|^2 \rangle$ averaged over production frames, with error bars indicating the standard deviation across frames. The solid line shows the two-step fit to the continuum membrane model including bending rigidity $\kappa$, surface tension $\Sigma$, and an additive noise floor $C$. Vertical lines indicate the high-$l$ and low-$l$ fitting windows used to separately determine $\kappa$ and $\Sigma$.

combined to define the mid-surface geometry,

$$r_{\text{mid}} = \frac{1}{2}\left(r_{\text{in}} + r_{\text{out}}\right), \qquad \mathbf{x}_{\text{mid}} = \mathbf{x}_{\text{COM}} + r_{\text{mid}}\,\hat{\mathbf{u}}_{\text{mid}}. \tag{72}$$

The inertia tensor of the mid-surface point cloud was computed about its center-of-mass, and its eigenvalues $\lambda_z \geq \lambda_y \geq \lambda_x$ were used to evaluate the relative shape anisotropy,

$$\kappa^2 = \frac{3}{2}\frac{\lambda_x^2 + \lambda_y^2 + \lambda_z^2}{(\lambda_x + \lambda_y + \lambda_z)^2} - \frac{1}{2} \tag{73}$$

The resulting $\kappa^2$ was reported as a function of timestep over the combined trajectory. The time evolution of the asphericity parameter $\kappa^2$ for all osmolyte concentrations considered in this work is shown in Fig. 5.

**Vesicle Volume Calculation**

The enclosed vesicle volume was computed directly from membrane bead configurations using a three-dimensional voxel reconstruction approach. For each frame, membrane bead coordinates were mapped into the primary simulation

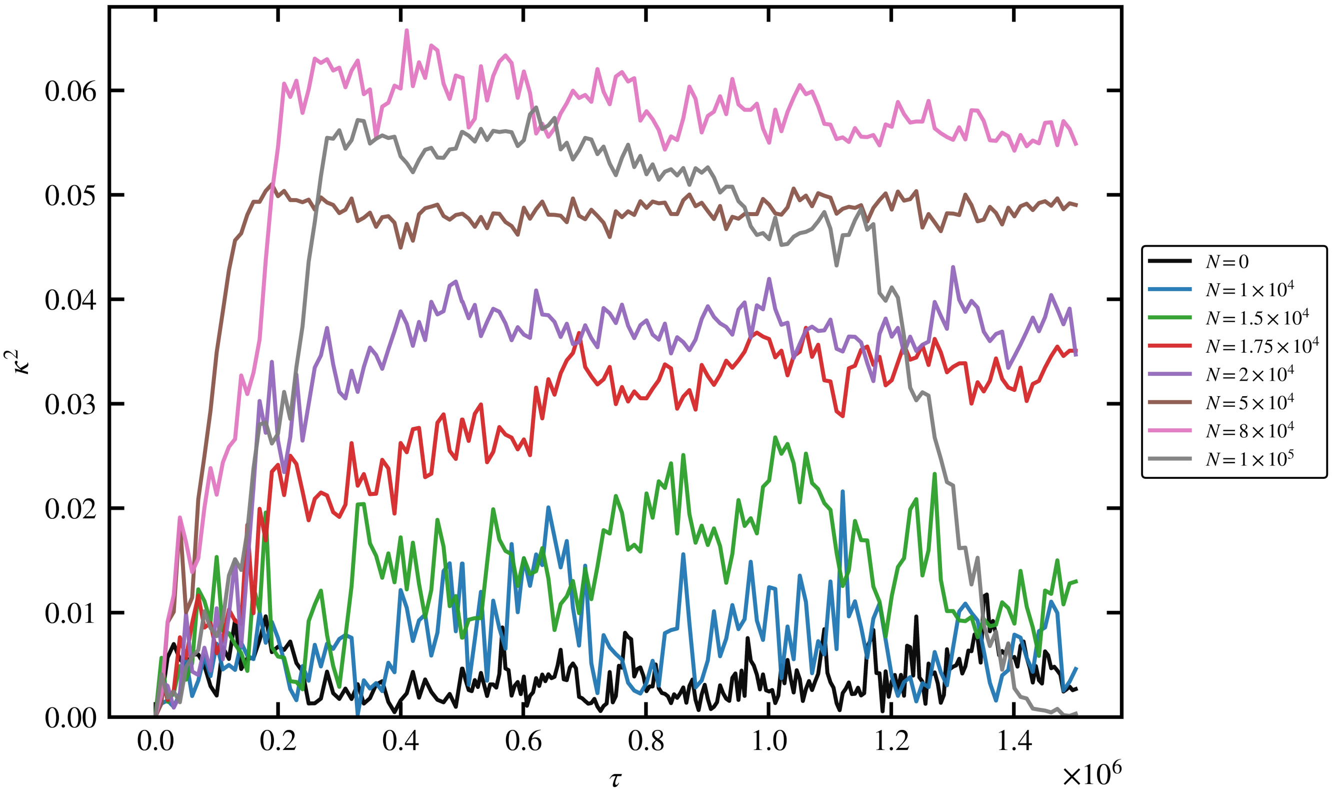


FIG. 5. Asphericity parameter $\kappa^2$ as a function of time for vesicle simulations with different osmolyte counts. We identify a critical osmolyte concentration $N_c \approx 1.75 \times 10^4$ using which we compute the critical pressure $\Delta p_c$.

box and recentered to ensure the vesicle was positioned away from periodic boundaries.

A cubic grid of spacing $\Delta x$ was superimposed over the simulation box, and voxels containing membrane beads were identified as part of the membrane surface. To ensure a closed surface representation, the membrane mask was slightly expanded to eliminate small gaps arising from discretization. A flood-fill algorithm was then applied starting from the box boundaries to identify all voxels connected to the exterior region. Voxels not connected to the exterior and not belonging to the membrane were classified as interior.

The vesicle volume was estimated as

$$V_{\text{ves}} = N_{\text{inside}} \, (\Delta x)^3, \tag{74}$$

where $N_{\text{inside}}$ is the number of interior voxels. Reported volumes correspond to time-averaged values computed over the production window after discarding initial equilibration frames. Statistical uncertainties were estimated from the standard error of the mean over the averaging window.


[1] W. Helfrich, Zeitschrift für Naturforschung C **28**, 693 (1973).
[2] H. J. Deuling and W. Helfrich, Biophysical Journal **16**, 861 (1976).
[3] M. Rubinstein and R. H. Colby, *Polymer Physics* (Oxford University Press, Oxford, 2003).
[4] F. Jülicher and U. Seifert, Physical Review E **49**, 4728 (1994).
[5] U. Seifert, K. Berndl, and R. Lipowsky, Physical Review A **44**, 1182 (1991).
[6] I. R. Cooke and M. Deserno, The Journal of Chemical Physics **123**, 10.1063/1.2135785 (2005).
[7] I. R. Cooke, K. Kremer, and M. Deserno, Phys. Rev. E **72**, 011506 (2005).
[8] S. Plimpton, Journal of Computational Physics **117**, 1 (1995).
[9] S. Biswas and B. Chakrabarti, Macromolecules **58**, 8091 (2025).
